# Phylogenetic tree inference from single-cell RNA sequencing data

**DOI:** 10.1101/2025.09.04.674140

**Authors:** Norio Zimmermann, Xiaoyu Sun, Joanna Hård, Jack Kuipers, Niko Beerenwinkel

## Abstract

Single-cell RNA sequencing technologies enable the large-scale measurement of gene expression profiles at the individual cell level to assess cellular diversity and function. In oncology, leveraging these single-cell transcriptomic data to reconstruct the phylogenetic relationships among cancer cells can provide insights into tumor evolution, metastasis formation, and the development of treatment resistance. However, phylogenetic inference from single-cell RNA sequencing is challenging due to sparse and noisy data and large dataset sizes. We present a novel tree inference method designed for such data that takes reference and alternative read counts of single-nucleotide variants and reconstructs a phylogenetic tree of the sequenced cells via maximum likelihood using a random-scan greedy search. To overcome local optima in the search, our algorithm alternates between two different tree representations: cell lineage trees, where cells are represented by nodes and mutations are attached to edges, and mutation trees, where mutation nodes encode the mutational events and cells are attached to them. Because a local optimum in one tree space generally does not correspond to a local optimum in the other space, we maximize the likelihood by switching between the two tree spaces until convergence is achieved in both. We demonstrate superior performance of our approach on simulated data with complex clonal architectures compared to existing methods. Furthermore, we show its applicability to cancer single-cell RNA sequencing data, which allows us to link evolutionary trajectories of cells to their gene expression profiles.

## 1 Introduction

Recent advancements in sequencing technologies have led to an increase in the availability of single-cell sequencing data and the development of new computational tools (5). In contrast to conventional methods such as bulk sequencing, single-cell sequencing technologies provide higher resolution for analyzing the heterogeneity of cell populations. In single-cell RNA sequencing (scRNA-seq), amplified complementary DNA (cDNA) obtained by reverse transcription of RNA is sequenced (20). Since the transcriptome is transcribed from the genome, scRNA-seq can indirectly provide genetic information about the expressed genomic regions, along with providing insights into the gene expression profiles of cells (5). It is therefore, in principle, also informative about the evolutionary relationships among cells. Phylogenetic inference aims to reconstruct these evolutionary histories from observed heritable traits, such as genetic variations.

For single-cell DNA sequencing data, which directly measures genetic variation in the DNA, various phylogenetic tree inference methods have been developed (7; 10; 11; 13; 14; 15; 25; 26; 27). However, its adoption remains limited due to high costs (16). More commonly used are bulk DNA sequencing, as well as scRNA-seq (16). scRNA-seq technologies include full-length protocols such as Smart-seq2 (20), and high-throughput 3’-end methods (32) as offered by 10x Genomics. Full-length methods typically offer greater sensitivity for gene detection, with lower technical noise, higher coverage and fewer dropout events compared to 3’-end protocols, which additionally suffer from 3’-end coverage biases (30; 8). Nonetheless, 3’-end methods are advantageous for detecting rare cell populations due to their higher cell throughput (30).

Most scRNA-seq-based phylogenetic tree reconstruction methods rely on single-nucleotide variants (SNVs), and therefore benefit from the aforementioned advantages of full-length technologies like Smart-seq2 compared to 3’-end methods. scRNA-seq tree inference methods include DENDRO (33), SClineager (18), PhylinSic (17), Canopy2 (31), and PhylEx (9). Both DENDRO and SClineager use alternative and reference allele read counts of transcribed SNVs as input. DENDRO then computes a pairwise genetic divergence matrix between cells and clusters the cells based on this matrix into clones (33). SClineager aims to overcome drop-out by using a Bayesian hierarchical model with Markov Chain Monte Carlo (MCMC) sampling to infer the true variant allele frequencies of the cells. These are then used to build clonal trees by hierarchical clustering methods (18). PhylinSic calls SNVs and infers a nucleotide sequence from the read counts of the mutated loci before tree inference. Then it uses BEAST2 (1) models to infer phylogenetic cell lineage trees (17). Both PhylEx and Canopy2 integrate bulk DNA data with scRNA-seq data to reconstruct clonal trees. Phylex (9) uses a tree-structured stick breaking process and Canopy2 (31) a Bayesian hierarchical model with MCMC sampling.

The existing methods have several limitations. SClineager and DENDRO rely on clustering cells into a limited number of clones, which may lack resolution and not accurately reflect the true phylogenetic structure. By calling genotypes from reference and alternative read counts before reconstructing the phylogeny, PhylinSic does not fully account for the read count information during tree inference. PhylEx’s and Canopy2’s use of bulk DNA data poses limitations when only scRNA-seq data is available. Another common problem of tree optimization methods that explore the tree space, such as PhylinSic, PhylEx, and Canopy2, is their tendency to become trapped in local optima, which might be difficult to escape from.

In this paper, we present **S**ingle-**C**ell **I**nference of **T**umor **E**volution from **RNA** sequencing data (SCITE-RNA), a tree inference method for scRNA-seq data. SCITE-RNA takes reference and alternative read counts of SNVs called from scRNA-seq data, selects potentially mutated loci, and reconstructs a phylogenetic tree of the sequenced cells. In the inference we greedily optimize over cell lineage and mutation trees, by alternating between these two tree spaces. This approach is fast and has the ability to escape local optima, because a simple tree move in the mutation space, such as reattaching a subtree at a new location, can lead to major changes in the structure of the cell lineage tree and vice versa. Using SCITE-RNA we address some of the limitations of existing approaches: (1) SCITE-RNA does not rely on additional data sources like bulk DNA sequencing, (2) it does not assume a certain number of clones in advance, (3) it specifically addresses the problem of getting trapped in local optima, and (4) it can estimate important parameters, such as the dropout probability, directly from the data.

We show superior performance of SCITE-RNA on simulated data for difficult tree reconstruction problems involving a large number of clones. Additionally, we demonstrate its ability to reconstruct plausible phylogenies on several real cancer scRNA-seq datasets.

## 2 Results

We first present a short overview of the SCITE-RNA method, followed by a comparative evaluation on both simulated and real scRNA-seq datasets. We benchmark our model against existing phylogenetic tree inference approaches for scRNA-seq data.

### 2.1 Method overview

SCITE-RNA (Fig. 1) takes as input reference and alternative read counts from SNVs across individual cells. For each cell-mutation pair, it computes a likelihood based on a model of scRNA-seq data involving the observed read counts, the inferred genotype and model parameters such as the dropout rate. Using these likelihoods, the algorithm obtains the likelihood of any given mutation or cell lineage tree. For cell lineage trees, nodes represent individual cells and the binary tree structure captures the evolutionary relationships between them. For mutation trees, nodes correspond to mutations, and the tree encodes the order in which mutations occurred. We do not model branch lengths for either tree type. SCITE-RNA employs a greedy optimization strategy that iteratively explores the tree spaces via random pruning and reattachment operations. Since this process is stochastic, the inferred tree and its likelihood can vary from run to run. After each optimization round, the tree is converted to its alternative representation, either from a cell lineage tree to a mutation tree or vice versa (Fig. 2). This iterative process of tree optimization and space switching is continued until the likelihood reaches an optimum in both spaces. Finally, based on the inferred genotypes obtained from the optimized tree, SCITE-RNA refines model parameters such as the dropout rate and overdispersion parameters. They influence the probability of observing specific reference and alternative read counts given a genotype and can vary across datasets or even across individual SNVs. The parameter optimization thus also enables the model to better account for RNA-specific phenomena, such as monoallelic expression and imbalanced biallelic expression of different genes. The updated parameters are then used in a second round of tree inference since the optimal tree structure may depend on the model parameters. The output of SCITE-RNA composes both the tumor single-cell phylogeny and the inferred genotypes (Fig. 1). A complete description of the methodology can be found in section 3.

**Fig. 1.**
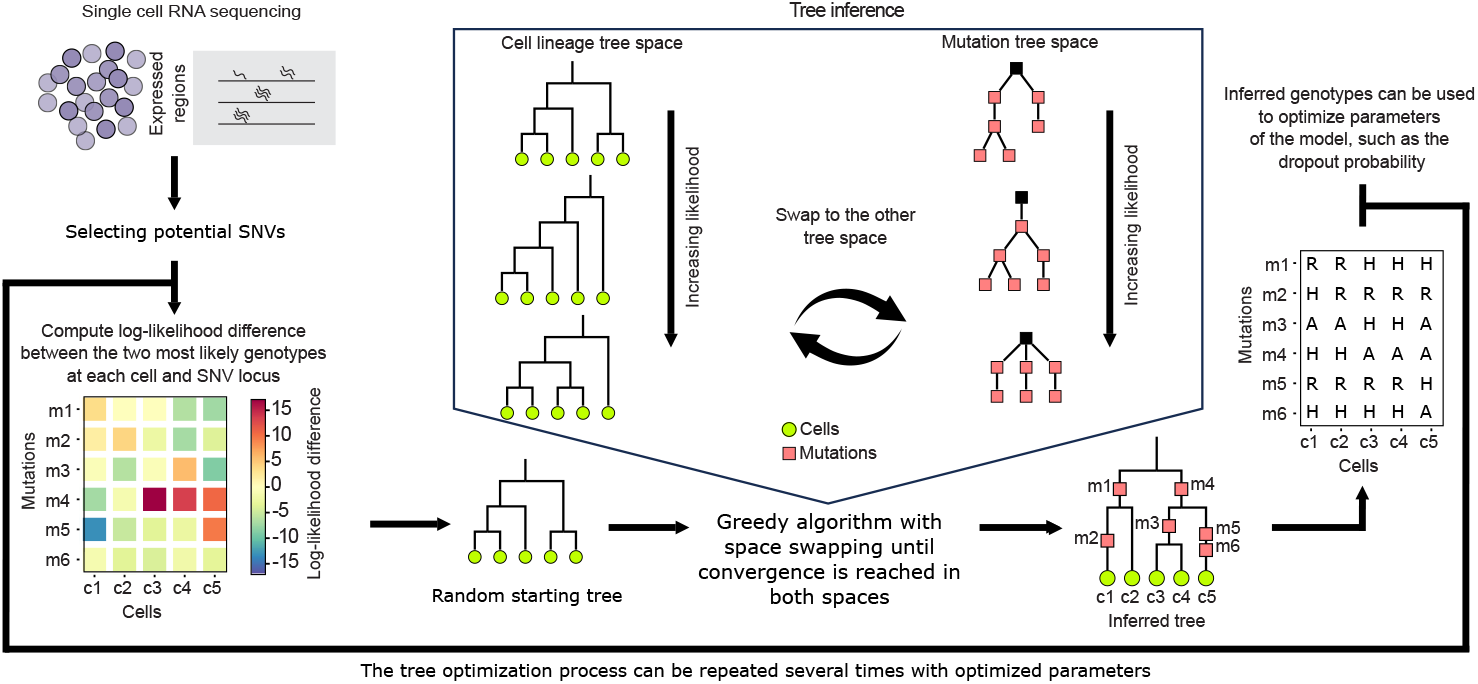
SCITE-RNA method overview. First, RNA of single cells is sequenced, followed by the selection of potentially mutated loci. We then obtain a likelihood for the read counts of each cell-mutation pair given the inferred genotypes and hence a joint likelihood for any given cell lineage tree or mutation tree. To find the maximum likelihood tree, we explore the tree space greedily with simple tree moves. Once a set of moves is completed, the tree is converted into the alternative space, where the exploration continues. This process is repeated until the tree hits a local optimum in both spaces. Using the genotypes inferred by the final tree, we optimize the model parameters to better fit a specific dataset and continue with the second round of tree optimization.

**Fig. 2.**
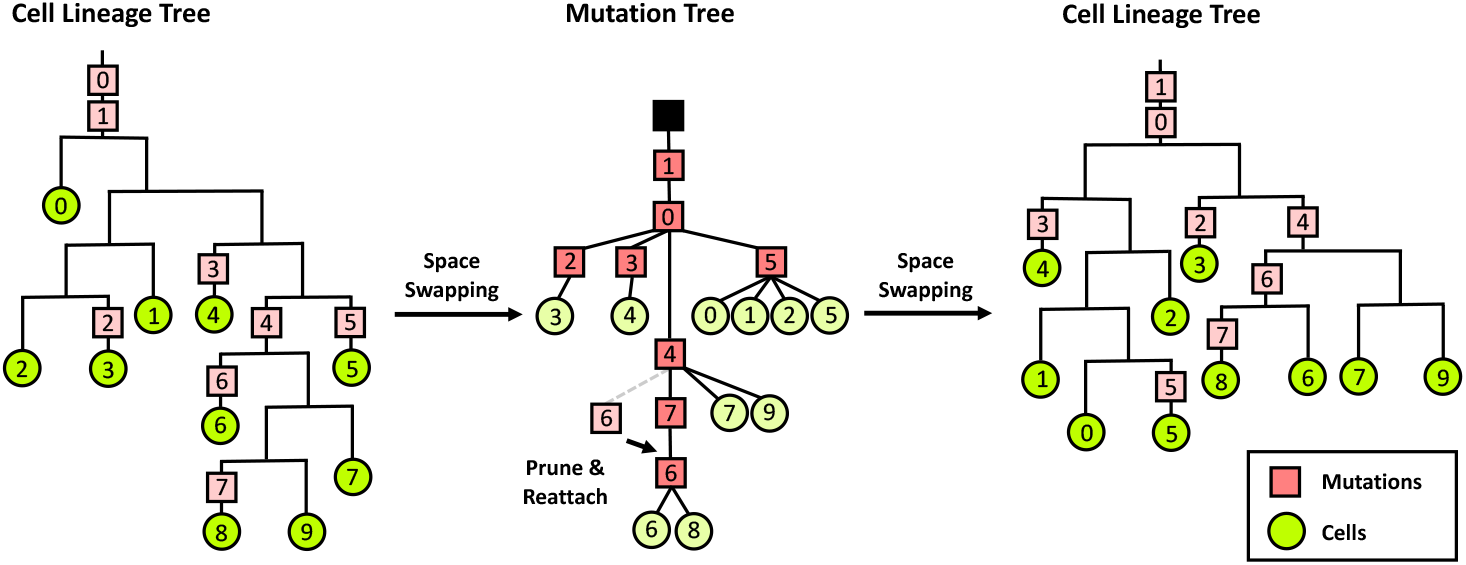
Swapping between tree spaces. The cell lineage tree on the left has been optimized by pruning and reattaching subtrees of cells. It is converted into a mutation tree representation (middle). Mutations placed on the same edge are assigned a random order during the tree space swapping. A simple move in the mutation tree space, for example, moving a single mutation (No. 6) to a new location, increases the likelihood of the tree. Swapping back to the cell lineage tree representation, the simple move in the mutation tree, combined with random sampling among equivalent tree conversions, has resulted in significant changes to the cell lineage tree.

### 2.2 Performance evaluation on simulated data

#### Comparison of tree space optimization strategies

To compare optimization in different tree spaces (Fig. 3), we conducted a simulation study using three distinct configurations of cells and SNVs: more SNVs (100 cells, 500 SNVs), balanced (500 cells, 500 SNVs), and more cells (500 cells, 100 SNVs). We simulate 100 random binary cell lineage trees with randomly placed mutations in each. Tree optimization is compared across four approaches: optimizing only in cell lineage tree space (c), only in mutation tree space (m), and alternating between the two, starting in either cell tree space (cm) or mutation tree space (mc).

**Fig. 3.**
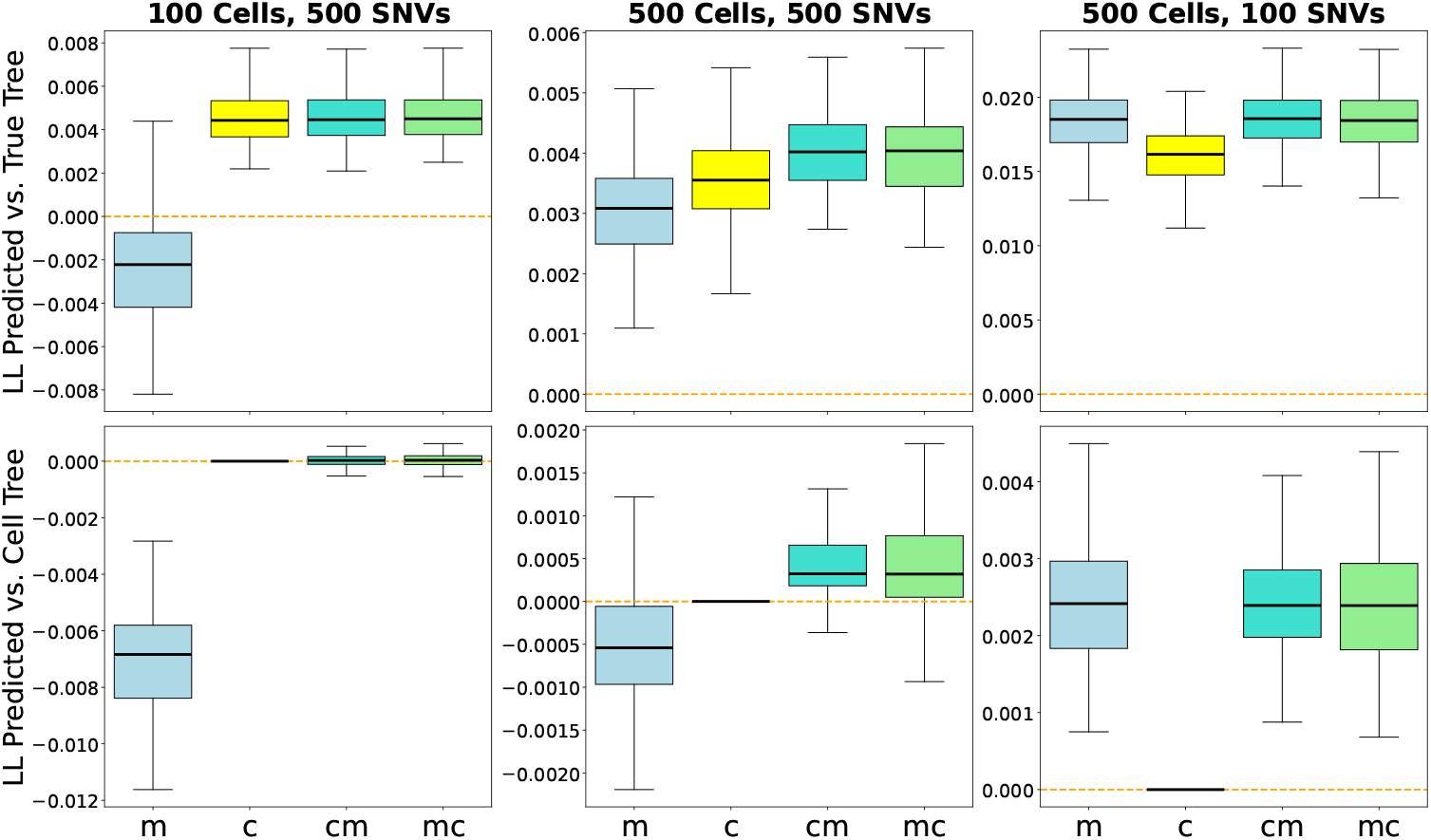
Comparison of tree optimization strategies across different configurations of cells and SNVs. The upper row shows the normalized log-likelihood (LL) difference between the optimized trees and the ground truth trees. Values above 0 indicate, that the optimized trees have a higher likelihood than the true trees. The lower row shows the normalized log-likelihoods after subtracting the log-likelihoods of the trees optimized solely in the cell lineage tree space. This highlights the differences in achieved optimal log-likelihood for individual runs. Tree optimization is compared across four approaches: optimizing only in cell lineage tree space (c), only in mutation tree space (m), and alternating between the two, starting with either cell tree space (cm) or mutation tree space (mc).

We compare the likelihood of the optimal trees to that of the ground truth trees, normalized by the number of cells and SNVs. Values greater than 0 indicate that the inferred trees have a higher likelihood than the ground truth trees, which is possible due to finite sample noise.

As shown in Figure 3 SCITE-RNA finds high-likelihood trees for all approaches, with the exception of solely optimizing in the mutation tree space (m) for larger numbers of SNVs. The benefit of switching between tree spaces becomes more pronounced when the number of cells is similar to the number of SNVs. When the number of SNVs is larger than the number of cells, optimizing the cell tree space is more effective due to its smaller size. The opposite, but less pronounced effect is observed, when the number of SNVs is smaller than the number of cells.

For an equal number of cells and SNVs, we observe that the likelihood, when switching between tree spaces, is usually higher compared to optimizing in a single tree space. When averaging across multiple runs, tree inference with space switching consistently performs at least as well as, and often superior to, the best results obtained by optimization in either tree space alone. This trend holds robustly across all tested combinations of cell and mutation counts.

#### Comparison with existing methods

We compare the performance of SCITE-RNA to DENDRO (33) and SClineager (18) on simulated data. Like SCITE-RNA, both of these methods work with reference and alternative allele counts derived from scRNA-seq data. The evaluation focuses on the accuracy of predicted genotypes and the similarity of the inferred tree structure to the true tree. We quantify these aspects by calculating the mean absolute error of the predicted variant allele frequencies and the Path Difference between the ground truth and predicted trees (adapted from (28); see Supplementary Section A.1 for further details). We simulate trees with 50 cells and 500 SNVs, along with a variable number of clones, representing cells with the same genotype (Fig. 4). We chose a relatively small number of cells and large number of SNVs, as this configuration is similar to the real datasets we explore below. We generate random cell lineage trees with 5, 10, 20, and 50 clones. For each number of clones, we simulate 100 trees with reference and alternative read counts for each cell–locus pair (see Supplementary Section A.1 for more details).

**Fig. 4.**
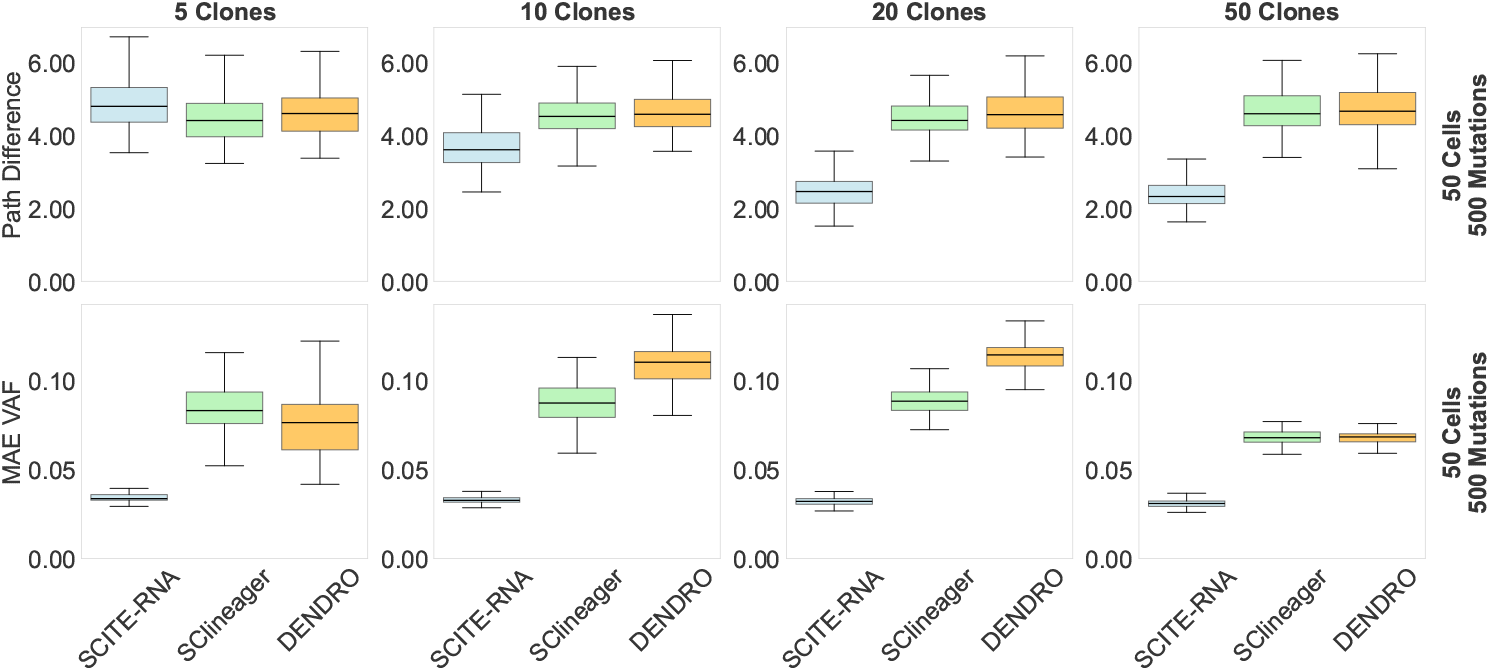
Comparison of SCITE-RNA to DENDRO and SClineager based on the Path Difference and the mean absolute error of the predicted variant allele frequencies (MAE VAF). The datasets simulate a variable number of clones with an increasing clonal complexity from left to right. SCITE-RNA outperforms the other methods in tree reconstruction for datasets with a higher number of clones and demonstrates superior accuracy in inferring true variant allele frequencies across all settings.

As shown in Figure 4 SCITE-RNA clearly and consistently outperforms the other methods in inferring the genotypes in all settings, while it reconstructs more accurate tree structures in most settings. The benefit of using SCITE-RNA for tree inference increases with increased clonal complexity. However, for very small numbers of clones, the Path Difference to the true tree is slightly smaller for DENDRO and SClineager. This is because SCITE-RNA assumes that SNVs/cells are independently placed on the tree to factorize the likelihood, an assumption that does not hold when only a few clones are present. Combined with the noisy nature of the data, this leads to overfitting in the SCITE-RNA trees, as our model does not impose any constraints on the number of clones.

As SClineager and DENDRO use clustering to generate the tree structure, we also provide a comparison of our method using the same hierarchical clustering approach based on the predicted variant allele frequencies (see Supplementary Section A.7). With clustering, SCITE-RNA outperforms the other methods even for small numbers of clones, in terms of both tree reconstruction and cell-to-clone assignment (Supplementary Fig. A.2). For more complex tree structures, however, SCITE-RNA without clustering achieves a larger improvement in performance over DENDRO and SClineager compared to the clustered approach (Supplementary Fig. A.1).

Another approach to reduce overfitting a particular dataset is to generate a consensus tree from a set of bootstrapped trees (see Supplementary Section A.7).

This also offers improved performance (Supplementary Fig. A.1), particularly at low clonal complexity, where it is comparable to clustering. At moderate and higher complexity, the consensus tree outperforms the clustered tree, though the improvement over standard SCITE-RNA is modest. The runtime, however, increases linearly with the number of bootstrap samples, as a tradeoff for the possible improvements in accuracy. We apply this approach on the real datasets as well, as it yields more stable results than individual runs (see Section 2.3).

Regarding runtime, SClineager requires by far the most computational time with more than 50 seconds for a dataset with 100 cells and 100 mutations (see Supplementary Fig. A.3). DENDRO is the fastest finishing in less than a second even for datasets with a hundred cells and mutations. On similar datasets it takes a few seconds to run SCITE-RNA with two optimization rounds. SCITE-RNA therefore offers the best overall performance, at a relatively low computational cost compared to models using MCMC such as SClineager.

### 2.3 Application to cancer data

We applied our model to two scRNA-seq datasets from multiple myeloma patients, with identifiers MM16 and MM34 (4). For MM16, 46 cells were sampled from a bone marrow biopsy at diagnosis and again six months later during a minimal residual disease state following chemotherapy. The MM34 dataset includes 127 cells from both the primary tumor and a metastatic site. After preprocessing, we retained 1,015 SNVs in MM16 and 1,528 SNVs in MM34 for phylogenetic tree inference (see Supplementary Section A.2). The primary sample was collected via bone marrow biopsy, while the metastatic sample was obtained from an ascites punctuation after two months of unsuccessful treatment (4; 24). After preprocessing and filtering the data (see Supplementary Section A.2), we ran two rounds of tree inference and parameter optimization, from which we use the results of the second round. More information on the parameter optimization can be found in Supplementary Section A.3. In addition we generated a consensus tree from 1,000 bootstrap samples to assess uncertainty in the tree structure (see Supplementary Section A.7).

In the MM16 dataset, our inferred trees (Fig. 5) show a partial separation between pre-treatment and post-treatment cells, with some post-treatment cells (in blue) appearing more closely related to the primary tumor cells (in red). This observation aligns with findings from the original study (4), which identified five post-treatment cells as surviving tumor cells based on copy number alterations. In contrast, DENDRO separates the two samples into two almost completely distinct branches, which does not align with five of the post-treatment cells indeed being tumor cells similar to the pre-treatment cells. SClineager predicts more intermixing of pre- and post-treatment cells but also does not reflect the expected five cells that should appear more closely related to the pre-treatment cells.

**Fig. 5.**
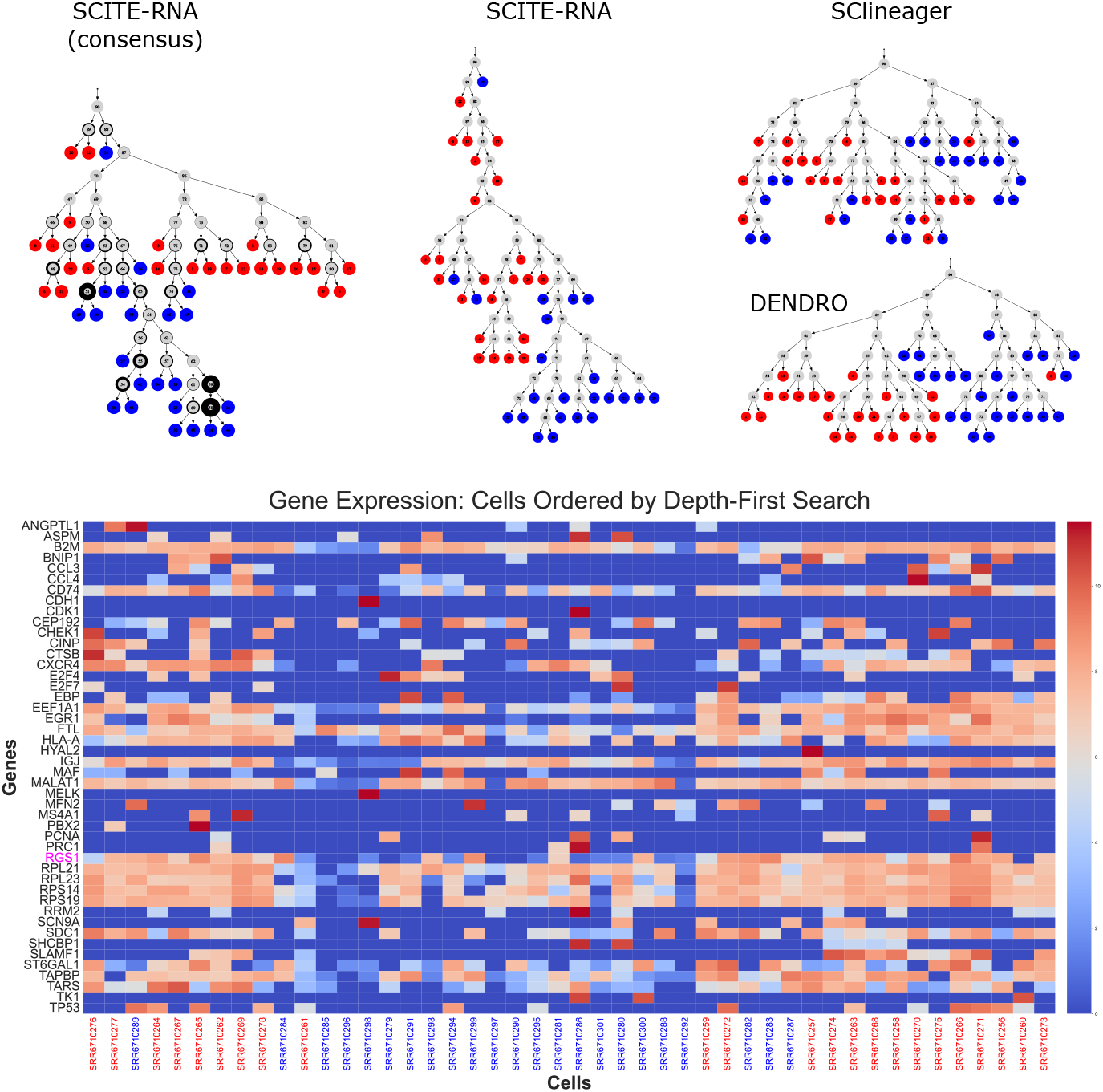
Application of SCITE-RNA, DENDRO and SClineager on multiple myeloma dataset MM16. A consensus tree was generated from 1,000 bootstrap samples. Red nodes represent primary tumor cells, blue nodes indicate post-treatment cells, and grey nodes correspond to unobserved ancestral states. The thickness of the black border around internal consensus tree nodes reflects the support for each split. Below the trees, we show the expression levels of selected genes, with columns ordered according to a depth-first traversal of the SCITE-RNA consensus tree. The gene RGS1 is highlighted in pink.

For patient MM34, our method identifies a clear split between primary and metastatic tumor cells, with metastasis-derived cells forming a distinct subtree (Fig. 6). The consensus tree topology supports a linear evolutionary model, consistent with the interpretation of the authors of the original study (4). This expected separation between primary and metastatic tumor cells is not captured as clearly in the tree inferred by DENDRO and even less so in the tree inferred by SClineager.

**Fig. 6.**
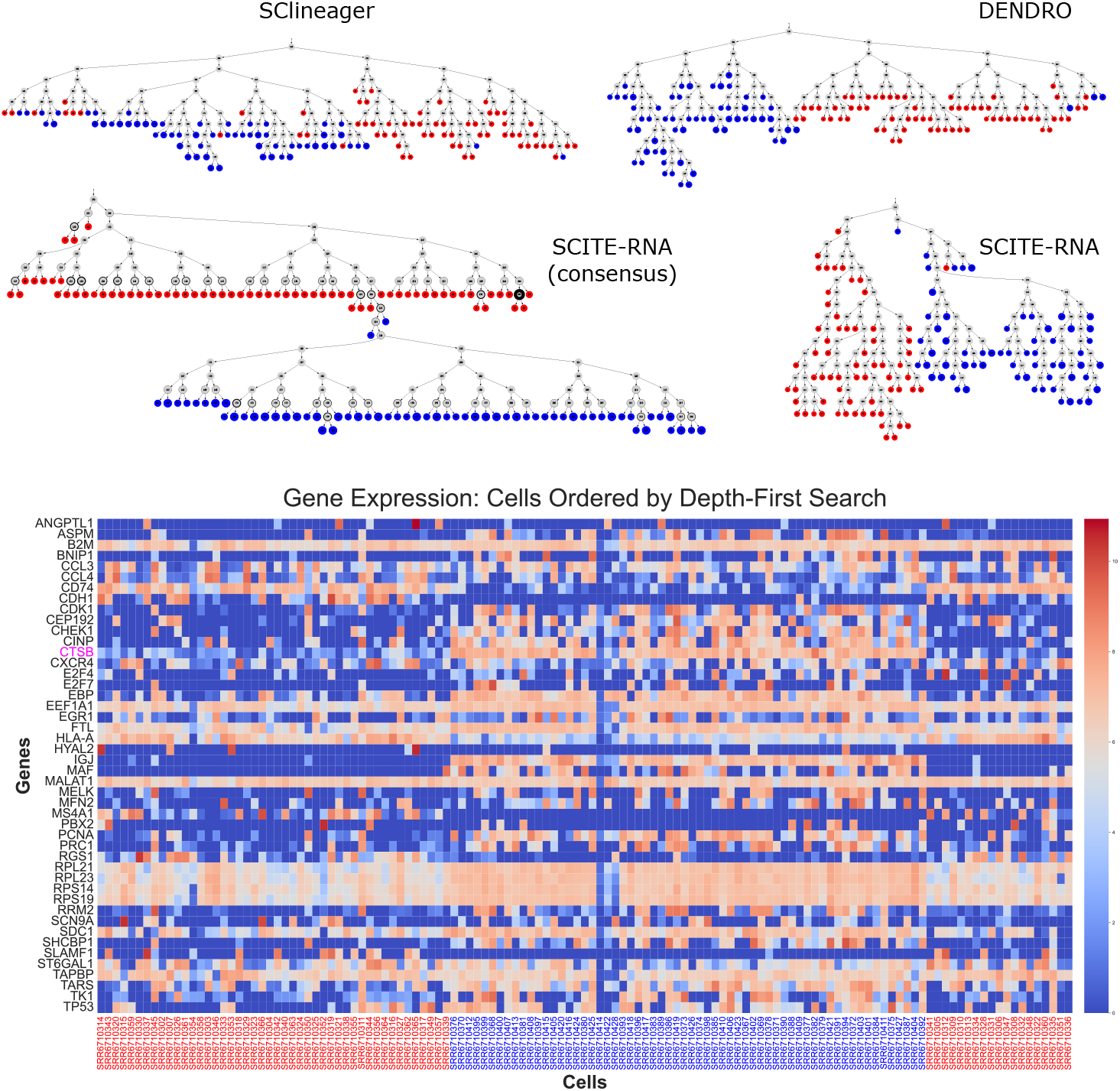
Application of SCITE-RNA, DENDRO and SClineager on multiple myeloma dataset MM34. A consensus tree was generated from 1,000 bootstrap samples. Red nodes correspond to primary tumor cells, while blue nodes indicate metastasis cells. Grey nodes represent unobserved internal ancestral states. The thickness of the black border around internal consensus tree nodes reflects the support for each split. Below the trees, we show the expression levels of selected genes, with columns ordered according to a depth-first traversal of the consensus SCITE-RNA tree. The gene CTSB is highlighted in pink.

In addition to the tree structure, scRNA-seq data enables us to explore gene expression patterns across the inferred phylogenies (Fig. 5 6). We visualize the expression of several genes highlighted by Fan et al. (4) for their potential relevance to multiple myeloma, ordering cells left to right via depth-first traversal of the SCITE-RNA tree. The expression patterns seem to reflect the branching structure of the tree, despite some cells originating from different samples. They also provide additional biological support: In MM16, for instance, the gene *RGS1* (regulator of G-protein signaling 1, highlighted in Fig. 5), which is associated with poor prognosis in multiple myeloma (23), is markedly more expressed in the branches closely related to the pre-treatment cells. In MM34, *CTSB* (Cathepsin B, highlighted in Fig. 6), whose overexpression is known to be associated with various cancers (19), shows substantially higher expression in the metastasis branch compared to primary tumor cells.

## 3 Methods

We first outline the model assumptions and then describe the tree inference procedure.

### 3.1 Model assumptions

The model is based on the following assumptions: (1) Each locus has exactly two copies in each cell. (2) Each locus has two possible alleles, called the reference and the alternative allele. (3) Mutations always occur at individual loci and convert one of the two copies into the other allele. (4) At most one mutation can occur at any given locus (infinite sites assumption (12)). (5) Observed read counts per cell and locus are independent given the genotypes.

These assumptions allow for our method to be computationally efficient. However, the model does not account for more complex forms of genetic variation, such as copy number alterations or structural variants. Additionally, the infinite sites assumption can be violated in cases of recurrent mutations or back-mutations, especially in larger datasets or over long timescales.

We opted for these simplifications because they not only enhance the computational efficiency and thus allow for the optimization of large trees, but also reduce the risk of overfitting to sparse and noisy single-cell RNA-seq data by lowering model complexity.

Under these assumptions, each cell contains either two reference alleles, two alternative alleles, or one reference allele and one alternative allele at any locus. These three genotypes are labeled **0** (reference-only), **1** (heterozygous), and **2** (alternative-only), respectively. Furthermore, there are only four types of transitions: **01** (i.e., **0** to **1**), **12, 10**, and **21**. A direct conversion between genotypes **0** and **2** is not possible because it would require two mutations.

SCITE-RNA takes candidate SNVs from scRNA-seq data as input in the form of a *N* × *M* matrix **D** of observations, where *N* is the number of cells and *M* the number of loci. An entry *D*_*ij*_ = (*r*_*ij*_, *a*_*ij*_) in **D** consists of the read count of the reference, *r*_*ij*_, and of the alternative allele, *a*_*ij*_, at locus *j* in cell *i*. As in (26), we treat the coverage *c*_*ij*_ = *r*_*ij*_ + *a*_*ij*_ as given and use beta-binomial distributions for *D*_*ij*_ when the genotype *G*_*ij*_ ∈ {**0, 1, 2}** is given.

For homozygous genotypes, the read counts are modeled as

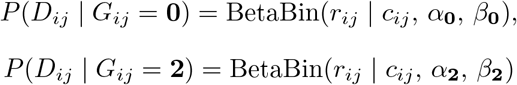

where *α* and *β* are parameterized by the expected variant allele frequency of the alternative allele *f* = *α/*(*α* + *β*) and overdispersion *ω* = *α* + *β*, which increases with decreasing variance. These parameters depend on the underlying genotype, as we detail later.

In the heterozygous case, we use a mixture of beta-binomial distributions to model allele-specific dropout. Specifically, we assume that when dropout occurs, the observed read counts follow the homozygous distribution of the remaining allele. Let *δ* denote the dropout probability. Then we set

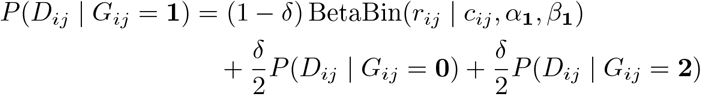

Since the observations per cell and locus are independent given the genotypes, the likelihood of **D** factorizes as the product over all cells and loci as

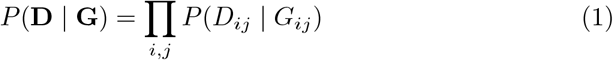

To estimate the parameters *α* and *β*, or equivalently *f* and *ω*, which are dependent on the genotype, we consider the case *G*_*ij*_ = **0**. Since in this case only reference alleles are present, all reads except the erroneous ones support that allele. Errors might arise during reverse transcription, PCR amplification, or during sequencing, and we assume these error processes affect both alleles symmetrically. Consequently, the expected variant allele frequency for the homozygous reference genotype should satisfy

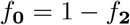

In the heterozygous case (*G*_*ij*_ = **1**), the two alleles are present in equal amounts, so their expected read counts are also equal, hence 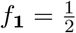.

To estimate the values of *ω*_**0**_ = *ω*_**2**_ and *ω*_**1**_, we note that in the heterozygous case, both biological and technical factors introduce additional variance to the expected variant allele frequency *f*. Reverse transcription may fail to capture one of the two alleles, and amplification and expression levels may differ between the two alleles. In such cases, the ratio between reference and alternative alleles is no longer 1:1 at sequencing time. This variability is accounted for by choosing a smaller value for *ω*_**1**_.

To infer the expected variant allele frequency *f*_**0**_, dropout *δ*, and the overdispersion parameters *ω*_**0**_ and *ω*_**1**_ from real scRNA-seq data, we adopt an iterative approach. In a first round, we perform tree inference using a predefined set of parameters. Based on the resulting genotype assignments, we optimize model parameters to better reflect the characteristics of the specific dataset. Since allelic expression can be imbalanced or even monoallelic in single cells, both *ω*_**1**_ and the dropout probability can vary across SNVs and consequently we optimize SNV-specific parameter values (see Supplementary Section A.3 for more details). These updated parameters of the expected variant allele frequency *f*_**0**_, dropout *δ*, and the overdispersion parameters *ω*_**0**_ and *ω*_**1**_ serve as refined parameters for a second round of tree inference. By default, we perform two rounds of alternating tree reconstruction and parameter estimation, with the goal to improve model fit to the data and the quality of the inferred trees, while keeping the computational costs low.

### 3.2 Tree inference

A cell phylogeny can be represented as a cell lineage tree, where mutations are placed on the edges to show where they occur, or as a mutation tree, illustrating the tumor’s mutational history, with cells linked to specific mutations (Fig. 2). Tracing back to the root then identifies all mutations present in a given cell. Given a tree in either form and the attached mutations or cells, we can deduce the corresponding genotype profile **G**, which allows us to obtain the likelihood of that tree by multiplying the likelihoods of the data given the genotype for each cell–locus pair using Eq. 1. This mapping is not injective, as the same genotype profile can result from different trees, but we can group trees that produce the same genotype profile into equivalence classes. Then there is a one-to-one correspondence between classes in the two tree spaces and it is easy to convert a tree into a random instance of the same class (which therefore has the same likelihood) in the other space. As an example in Figure 2 consider the cell lineage tree with 2 mutation events occurring on the same edge at the root. If we convert this cell tree into a mutation tree, any permutation of these mutations (e.g. (0,1) or (1,0)) is valid and thus one mutation order is randomly chosen.

Before tree inference, potentially mutated loci are selected, and the most likely genotypes before and after the mutation are determined (see Supplementary Section A.4 for details). This allows us to compute the log-likelihood differences between those two genotypes for each cell–locus pair.

The goal of the tree inference algorithm is to find the tree with the highest likelihood. Due to the enormous sizes of the tree spaces, it is impractical to test every possible tree. Instead, the algorithm starts with a random tree and gradually optimizes it using a greedy approach. At each step, the algorithm is given a starting tree from which it explores the local tree neighborhood by performing tree operations (Fig. 1). During cell lineage tree optimization, nodes are selected in a random order. Then the respective subtrees consisting of the chosen node and its descendants are pruned and reattached at their optimal location. If there are several positions with equal likelihood, one is chosen randomly. This step is repeated until every node, including internal nodes, has been processed.

The same standard pruning and reattaching of subtrees is performed for mutation trees. Since they often contain linear chains of mutations, which are harder to optimize by switching subtrees, we added an optional second move for smaller mutation trees. In this move, individual mutations with only a single child are pruned and re-inserted at their optimal position. This helps optimally rearranging linear chains of mutations. For cancer datasets with several thousand potential SNVs, we focused on pruning and reattaching of subtrees to keep computational costs low.

For each tree, the algorithm calculates the highest likelihood achievable by attaching mutations to edges (for cell lineage trees) or cells to nodes (for mutation trees). Since the observations per cell and locus are assumed to be independent, we only need to determine the maximum-likelihood attachment location of each individual cell or mutation separately. We use established tree traversals (e.g. (7)) to lower complexity, which leads to our method scaling well with the number of cells and mutations (see Supplementary Section A.6 for details).

It is very common for greedy algorithms to get stuck in local optima. However, we optimize the tree by alternating between two different tree spaces, and a local optimum in one space does not necessarily correspond to a local optimum in the other space. Hence, the tree space swapping is a way to potentially escape from local optima (Fig. 3). The process of converting the tree into one possible representation in the other space and continuing the exploration there repeats until the likelihood in both spaces stops improving.

## 4 Discussion

We have proposed SCITE-RNA for inferring phylogenies from scRNA-seq data using SNVs and demonstrated that it outperforms existing approaches on simulated datasets (Fig. 4). Two alternative methods, DENDRO (33) and SClineager (18), focus on identifying a small number of cell clones and may miss more fine-grained biological heterogeneity that SCITE-RNA can capture.

To further validate the method, we applied SCITE-RNA to two multiple myeloma datasets. The inferred phylogenies demonstrated more biologically plausible evolutionary relationships among cells from different samples compared to those generated by DENDRO and SClineager. Additionally, the use of scRNA-seq data allows for the mapping of gene expression values onto the tree structure. This not only provides support for the inferred trees but also potentially enables the study of gene expression and cellular functions at a clonal level.

Inferring trees with hundreds or even thousands of cells and mutations requires a fast tree optimization algorithm. To address this, we employ random-scan greedy search and space-swapping, where moves in each space are easy to compute efficiently, even though they might be equivalent to complex and computationally expensive moves in the other space (Fig. 2). Our results show that alternating between cell and mutation tree spaces yields superior results, particularly when both mutation tree optimization and cell lineage optimization are similarly effective (Fig. 3). Even if optimization in one of the two spaces is suboptimal, jointly optimizing across both spaces is, generally, at least as effective as the better-performing individual space. Consequently, tree space switching is more robust regardless of the ratio of number of cells to SNVs. The tree likelihood of the inferred trees using tree space swapping was generally higher compared to the ground truth tree (Fig. 3). This highlights that the algorithm is achieving its objective of optimizing the trees well.

Nevertheless, overfitting remains a concern, particularly with noisy data. One cause is that placing each mutation at its individually optimal position in the tree may not yield a globally optimal solution. To mitigate this risk, an approach could be to penalize the number of edges with mutations for cell lineage trees and attachment points of cells in the mutation tree space to reduce the number of inferred clones. Alternatively, bootstrapping the mutations and then generating a consensus tree can reduce overfitting and allow the assessment of uncertainty in the tree structure. However, the runtime increases linearly with the number of bootstrap samples. A faster option to simplify trees is to use hierarchical clustering to group the cells into a smaller number of clones based on the predicted genotypes. We saw that both clustering and bootstrapping can offer improvements in performance for trees with few clones, and bootstrapping also for more complex phylogenies.

Another source of error involves the estimation of model parameters, which require accurate genotype predictions. Variant allele frequencies that are very low or high are more likely to be assigned homozygous genotypes, as these maximize the likelihood for individual cases. However, in the presence of allelic dropout, a notable proportion of these could be heterozygous, leading to an overestimation of homozygous genotypes by shifting the inferred mutation location in the cell lineage tree. Conversely, real datasets may contain loci with both reference and alternative genotypes due to multiple mutations affecting the same locus, violating our model assumptions. As the model only compares the two most likely genotypes, some cells with two mutations might be inferred as heterozygous, potentially overestimating the number of heterozygous genotypes. Furthermore, our method does not incorporate copy number alterations or gene expression in the tree optimization itself, both of which might also be obtained from scRNA-seq data, and offer interesting directions for further phylogenetic developments.

Despite these limitations, alternating between tree spaces for optimization is efficient and could be extended to other modalities, such as scDNA-seq data. The combination of fast optimization and data-driven estimation of model parameters, enables the use of scRNA-seq data not only for gene expression analysis but also for reconstructing the evolutionary history of cells, allowing us to augment expression-based analyses with phylogenetic information.

## Data availability

The multiple myeloma datasets are publicly available from the NCBI Gene-Expression Omnibus (GEO) under accession number GSE110499. The code is published on GitHub https://github.com/cbg-ethz/SCITE-RNA.

## Author Contributions

X.S., N.Z. and J.K. developed the method and contributed to data analysis. J.H. contributed to data analysis. J.K. and N.B. supervised the project. All authors contributed to the manuscript preparation.

## Acknowledgments

Part of this work was funded by the ERC Horizon 2020 program (project OLISSIPO, No. 951970) and the Swiss National Science Foundation (project No. 310030_179518 and 320030_236168).

## Supplementary Figures

**Supp. Fig. A.1.**
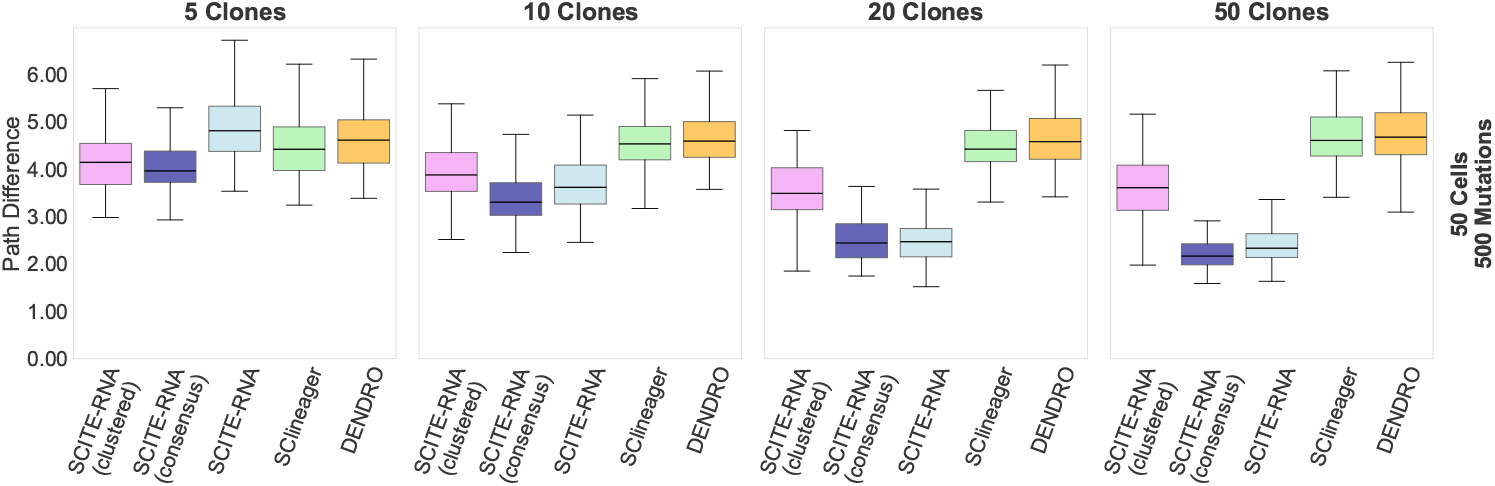
Comparison of SCITE-RNA, SCITE-RNA with subsequent clustering on predicted variant allele frequencies, SCITE-RNA with consensus tree generation from 1,000 bootstrapped trees, DENDRO, and SClineager using the Path Difference metric across simulated datasets. The datasets simulate a variable number of clones with an increasing clonal complexity from left to right.

**Supp. Fig. A.2.**
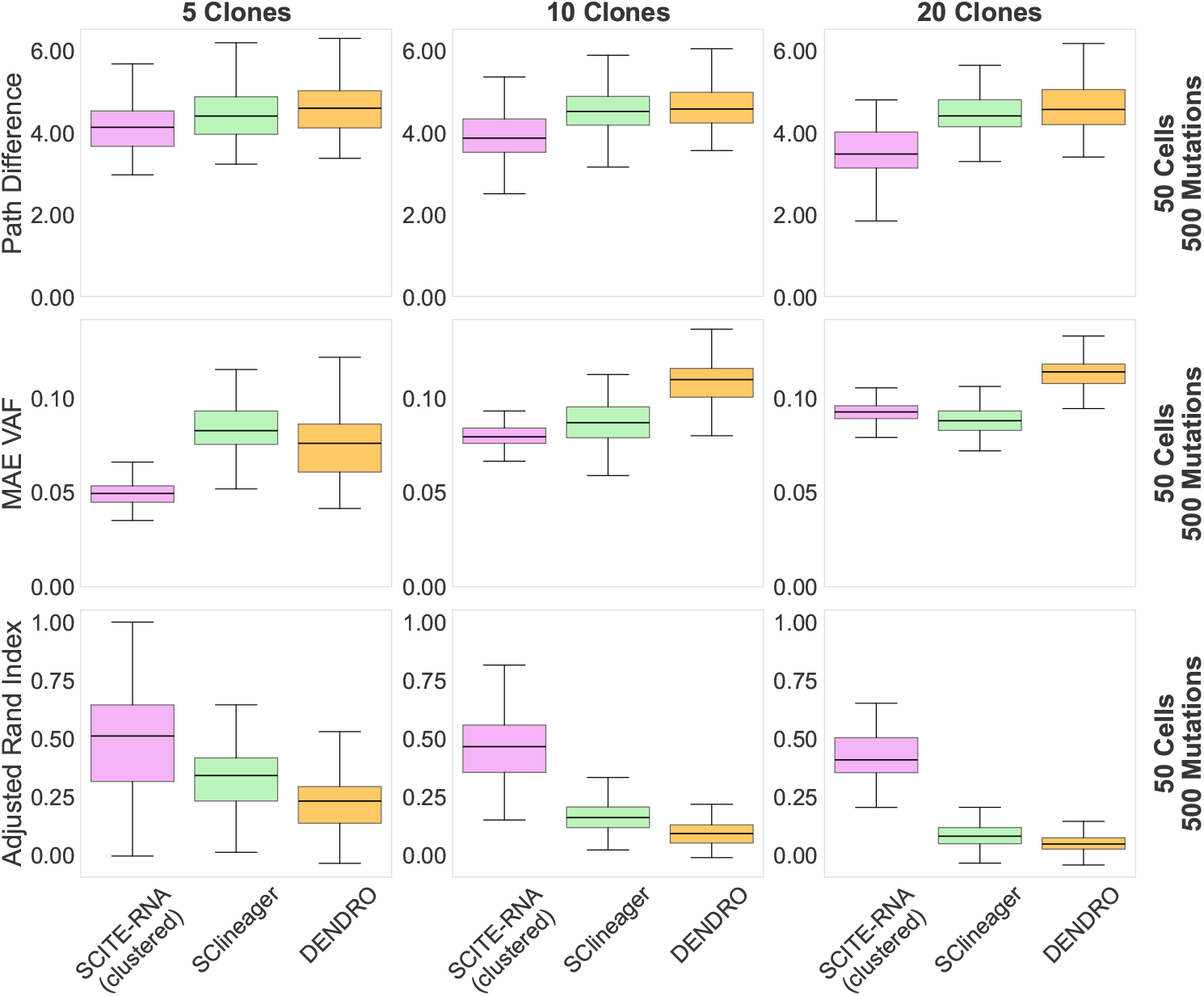
Comparison of SCITE-RNA with subsequent clustering on predicted variant allele frequencies, DENDRO and SClineager based on the Path Difference metric, the mean absolute error of the predicted variant allele frequencies (MAE VAF) and the Adjusted Rand Index. The datasets simulate a variable number of clones with an increasing clonal complexity from left to right.

**Supp. Fig. A.3.**
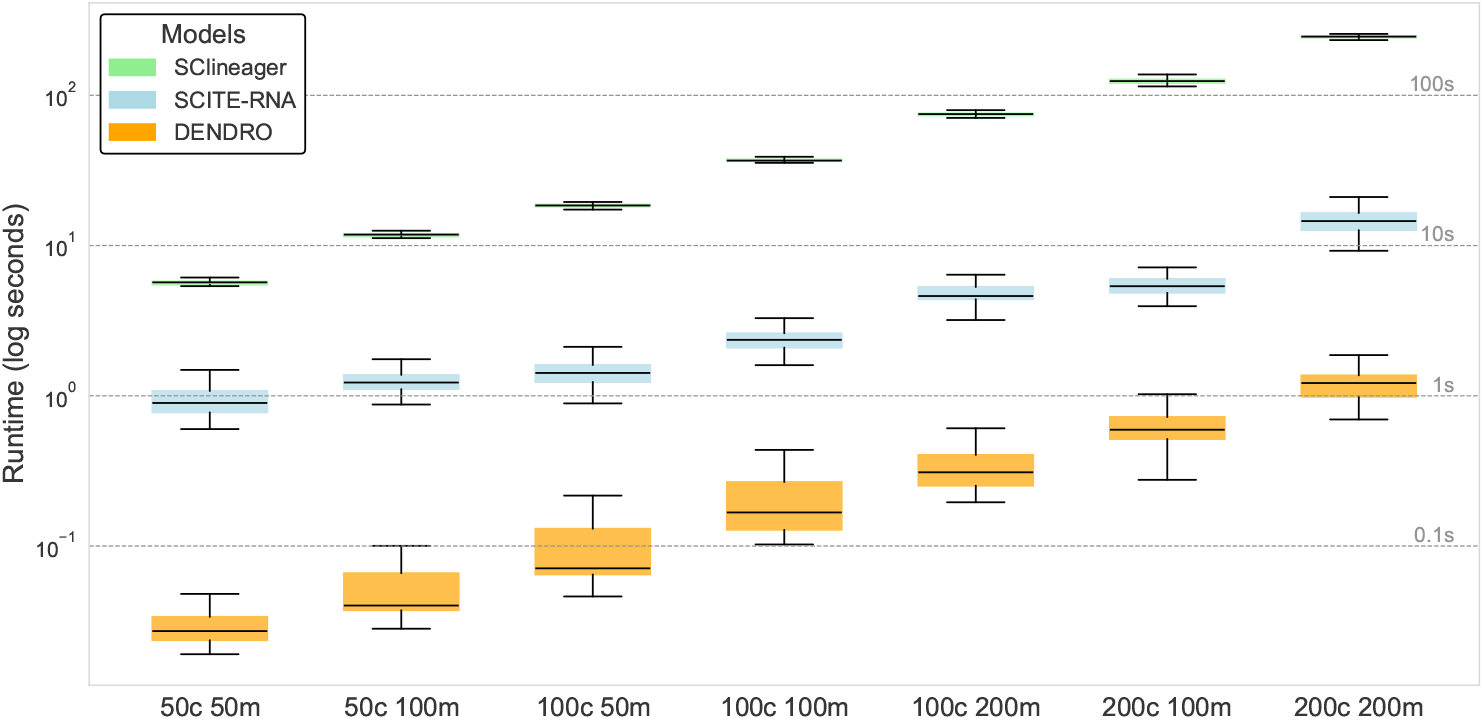
Comparison of runtimes between SCITE-RNA, SClineager and DENDRO for 100 simulated datasets with various numbers of cells and mutations. The runtime is plotted in seconds (log scale).

## A Appendix

### A.1 Simulation and Evaluation

The SCITE-RNA model and algorithms described above are implemented in Python and translated to C++ for improved performance. To simulate scRNA-seq data, first a random cell lineage tree is constructed. Mutations are either randomly attached or confined to fewer edges to control the number of clones. Here clones describe groups of cells with the same genotype. For each cell and locus, we sample sequencing coverage from a zero-inflated negative binomial distribution with mean *µ* = 60, zero-inflation probability *π* = 0.39, and dispersion *θ* = 0.17. The parameters of the distribution are fitted to match the cancer scRNA dataset MM34. The coverage and genotype are used to sample the read counts of the reference and alternative alleles from the beta-binomial distributions described in Section 3.1.

We compare our method, SCITE-RNA, with DENDRO (33) and SClineager (18). SClineager is run for 2000 MCMC iterations and outputs predicted variant allele frequencies. These are subsequently clustered using hierarchical clustering to assign cells to clones and infer a cell lineage tree. Similarly, DENDRO constructs a pairwise genetic divergence matrix between cells and applies hierarchical clustering with Ward’s method to assign clones and generate a tree (33).

Given that the genotypes in a simulated dataset are known, they can be compared with the predicted genotypes inferred from the tree. To assess the quality of these predictions, we use the mean absolute error of the predicted variant allele frequencies. The frequencies assigned to reference, heterozygous, and alternative genotypes are 0, 0.5, and 1, respectively. The variant allele frequencies predicted by SClineager and DENDRO are rounded to the nearest of those values before calculating the mean absolute error. Since DENDRO does not directly provide predicted variant allele frequencies, we fix the number of clones to the ground truth value and average the observed frequencies per clone. This approach could potentially favor DENDRO, as it typically relies on heuristics, such as inspecting the intra-cluster divergence curve, to determine the optimal number of clones (33).

To evaluate the accuracy of the tree topology, we use the Path Difference (adapted from (28)) between pairs of cells in the predicted and ground truth cell lineage trees. This metric counts the number of edges separating each cell pair in both trees and calculates the average absolute difference across all pairs. The original publication used the Euclidean distance, which is more dependent on the tree size and thus harder to compare across various tree sizes.

### A.2 Data processing

We downloaded the multiple myeloma datasets MM16 and MM34 from the NCBI Gene-Expression Omnibus (GEO) under accession number GSE110499. MM16 contains data from 23 pre-treatment and 23 post-treatment cells. MM34 consists of 65 cells sampled from the primary tumor and 62 from a metastasis.

**Supp. Fig. A.4.**
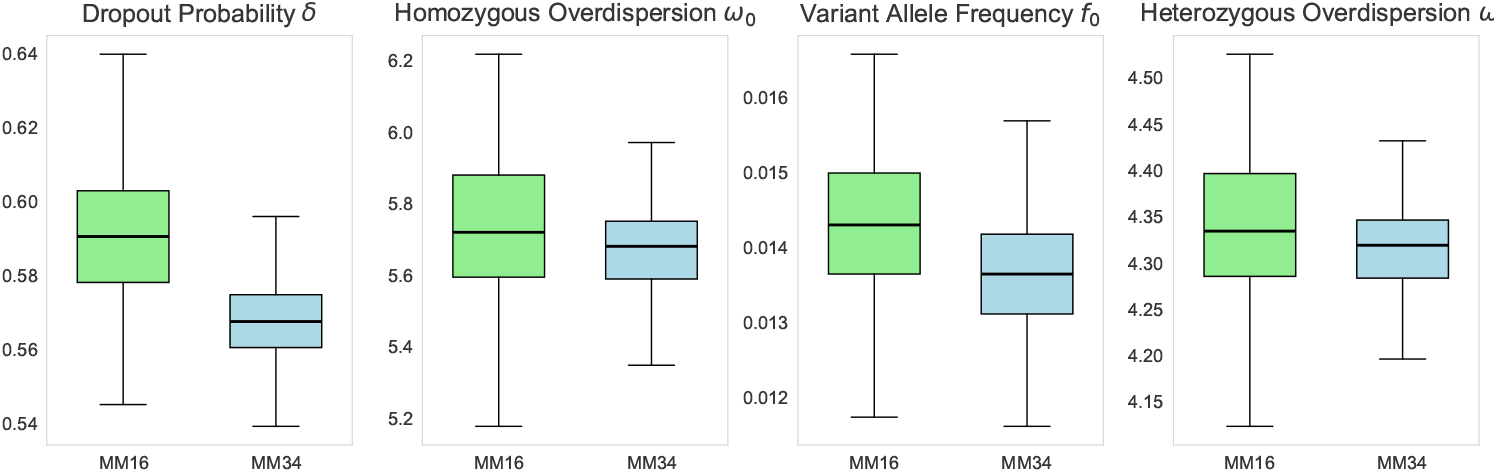
Estimated dropout *δ*, expected variant allele frequency *f*_**0**_, homozygous overdispersion *ω*_**0**_ and heterozygous overdispersion *ω*_**1**_ parameters over all loci using the inferred genotypes of 1000 bootstrap samples.

The samples were prepared using the C1 Single-Cell Auto Prep System (Fluidigm) and the SMARTer Ultra Low RNA Kit (Clontech) for cDNA synthesis and amplification. In addition, the HiSeq2500 (Illumina) was used for sequencing in 100-bp paired-end mode.

We applied STAR aligner (3) to map the reads to the human reference genome GRCh38. Following the approach outlined in Liu et al. (17), we combined the reads into a pseudo-bulk file and called variants using the GATK HaplotypeCaller (21). We only kept SNVs, i.e. no variants affecting larger regions of the genome and no multi-allelic sites. To reduce noise, we excluded loci with maximum coverage below 10, total coverage under 100, fewer than 5 alternative reads, or variant allele frequencies below 2%. To avoid very sparse data, we retained only loci with read counts measured in at least 25% of the cells. Additionally, if two loci were within 10 base pairs of each other, they were both removed, to prevent regions with numerous SNVs, potentially due to technical noise, from being over-represented in the dataset. The number of remaining potential SNVs after initial preprocessing was 1,178 for MM16 and 2,092 for MM34.

Using the mutation filtering step (see Supplementary Section A.4), we retained SNVs with a posterior probability greater than 5% of affecting at least two, but not all, cells in the tree. This conservative filtering criterion resulted in the selection of 1,015 SNVs for MM16 and 1,528 for MM34.

### A.3 Parameter estimation

As initial parameters, we used a dropout probability *δ* of 20%, an expected variant allele frequency *f*_**0**_ of 5%, a homozygous overdispersion *ω*_**0**_ of 10, and a heterozygous overdispersion *ω*_**1**_ of 6. Due to limited prior knowledge, we aimed to refine these values by learning them from the data. To achieve this we used the bounded limited-memory BFGS (Broyden–Fletcher–Goldfarb–Shanno) algorithm (2) to optimize the parameters by maximizing the likelihood of the observed read counts, given the inferred genotypes across all SNVs and cells. For those loci where at least five cells are inferred to be heterozygous and have coverage greater than 10, we defined SNV specific dropout and overdispersion parameters. For loci with fewer heterozygous cells and thus likely not enough data to infer better parameter estimates, we keep the globally inferred dropout probability and overdispersion *ω*_**1**_. Bootstrapping of mutations before tree inference, allows us to estimate the uncertainty in the parameter estimates. We observe that for the real cancer datasets the estimated dropout probabilities are relatively high, exceeding 50%. The overdispersion parameters are small, indicating substantial variance in the allele frequencies. The expected variant allele frequencies *f*_**0**_ are low, with a mean around 1.5% (Fig. A.4). The high dropout and low overdispersion parameter estimates could be explained by monoallelic expression and imbalanced allelic expression.

**Supp. Fig. A.5.**
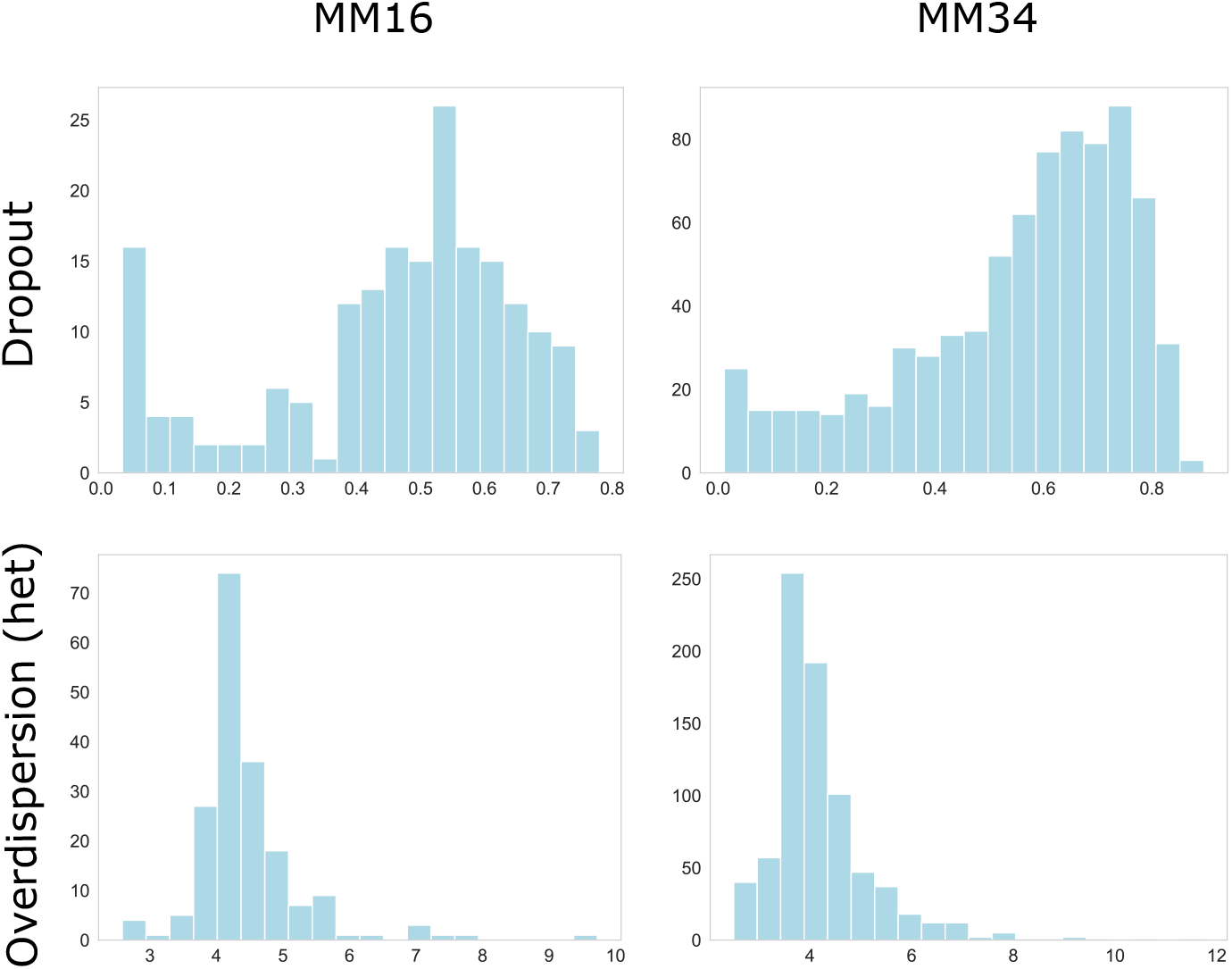
Estimated dropout and heterozygous overdispersion for individual loci using the inferred genotypes.

Additionally, we examine the distribution of dropout rates and heterozygous overdispersion values *ω*_**1**_ for individual SNVs. In this analysis, we compute the mean parameter estimates across 1000 bootstrap samples. Consequently, the resulting distribution reflects the variability of these parameters across different loci (Fig. A.5). We observe that estimates of the heterozygous overdispersion parameter (*ω*_**1**_) are centered around a value of about 3 to 4 across the datasets, with a small number of loci showing substantially higher overdispersion values. Dropout probabilities tend to be generally high, which can be expected for scRNA-seq data. However, each dataset also contains a notable subset of loci for which the estimated dropout probability is close to zero.

### A.4 Selection of mutated loci

To select loci that are informative about the structure of the phylogenetic tree and to infer the most likely genotypes at these loci, we proceed as follows.

First, suppose we know which cells are affected by a certain mutation, then we can rule out some of the possible binary cell lineage tree structures *T*_*N*_ with *N* leaves, because the correct tree must contain a subtree whose leaves are exactly those affected cells. Consequently, a mutation provides information about the tree structure only when it affects at least two cells and at most *N* − 1 cells, and we therefore aim to only maintain those mutations.

For each locus, we want to obtain the posterior probability that it contains an informative mutation. We define a prior for the mutation type and the number of affected cells. We assume equal prior probabilities for all genotype transitions (**01, 10, 21, 12**) and for the three genotypes (**0, 1** and **2**).

The prior for the number of affected cells is adopted from (26), where it is calculated by assuming all binary tree structures and mutation placements to be equally probable. We denote by *X* the wildtype and *Y* the genotype after mutation. As a result, the prior that *k* out of *N* cells are affected is

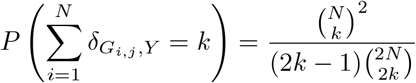

where *δ* is the Kronecker delta.

We obtain the overall mutation prior probability (i.e. the joint prior of the genotype transition and the number of affected cells) by multiplying the prior of the genotype transition with the prior of the number of affected cells.

To derive the posterior, we use that if a locus *j* is not mutated, all cells must have the same genotype *X* at *j*, and the joint likelihood of the corresponding observations **D**_∗*j*_ := {*D*_1*j*_, …, *D*_*Nj*_} is given by

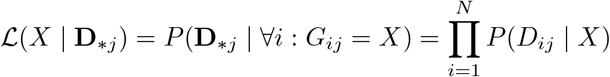

Otherwise, *j* is mutated with the genotype *Y* after mutation. We can calculate the likelihood that exactly *k* cells are affected (i.e., that *N* −*k* cells have genotype *X* and *k* cells have genotype *Y*) for any pair *XY*. The approach is detailed in Supplementary Section A.5.

Finally, the likelihood and prior can be multiplied and normalized to get the posterior probability of each mutation type *XY* ∈ mt := {**01, 10, 21, 12}** and number *k* of affected cells. Then we can marginalize over both the four genotype transitions and all the *k* values that are considered informative to get the posterior probability of the event *I*_*j*_ that locus *j* contains an informative mutation.

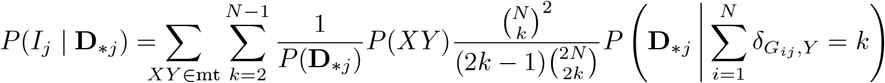

There are multiple ways to select loci based on the posterior, for example by choosing the *n* loci with the highest posterior or by setting a fixed cutoff value. For tree inference, we select the two most likely genotypes per locus. This allows us to pre-calculate the likelihood ratio for each cell/locus having genotype X vs genotype Y based on the read counts. We select the most likely genotypes based on the highest posterior of the four genotype transitions **01, 10, 21, 12**.

Selecting mutations based on the likelihood of the read counts for each cell individually (see Supplementary Section A.5) may be suboptimal in the case of high dropout rates or generally if there is too much overlap between the read count distributions of the genotypes. Cells with dropout of an allele usually have very high or low variant allele frequencies. Individually, these extreme variant allele frequencies are most likely explained by homozygous genotypes, while globally across all cells, you can expect a fraction of those cells to be heterozygous. Consequently, with our approach we are mostly able to filter out loci with all reference or all alternative genotypes, but would likely include loci with all heterozygous genotypes in the tree inference.

### A.5 Likelihood for the number of affected cells

We denote by *X* the wildtype and *Y* the genotype after mutation. For any *XY* combination of genotypes, we can use a dynamic programming approach to calculate the likelihood that exactly *k* cells are affected (i.e. that *N* − *k* cells have genotype *X* and *k* cells have genotype *Y*). We define *l*(*n, k*) as the likelihood, that k out of the first n cells are affected:

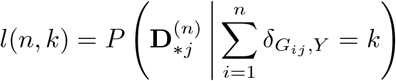

where 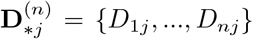 and *δ* is the Kronecker delta. When *n* = *k* = 0, the likelihood is *l*(0, 0) = 1, because there is only one possible 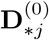, which is the empty set. When *n* ≥ 1, *l*(*n, k*) can be calculated recursively from *l*(*n* − 1, *k*) and *l*(*n* − 1, *k* − 1). This is achieved by first rewriting *l*(*n, k*) as a marginalization over all possible genotype combinations in the first *n* cells, with the condition that exactly k are affected:

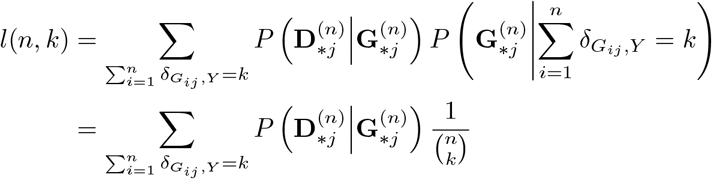

Where 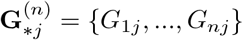. For convenience, we define:

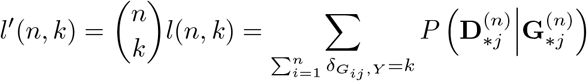

Next, we can divide the genotype combinations into two sets, depending on whether the *n*-th cell is mutated:

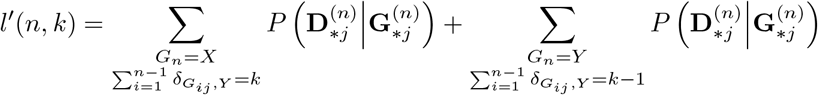

Note that in the first summation above, *P* (*D*_*nj*_|*X*) is a common factor and so is *P* (*D*_*nj*_|*Y*) in the second summation. Furthermore, if we take them out of the summations, the remaining parts become *l*^*′*^(*n −* 1, *k*) and *l*^*′*^(*n −* 1, *k −* 1) respectively. Thus, we have:

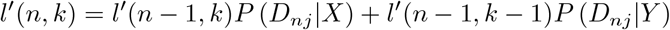

and consequently:

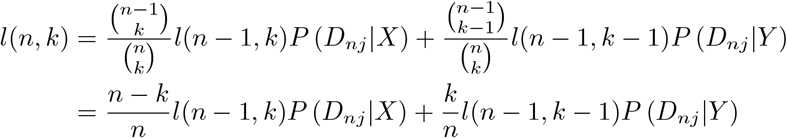

which allows us to eventually obtain *l*(*N, k*) for all numbers of affected cells *k*. If *l*(*n −* 1, *k*) or *l*(*n −* 1, *k −* 1) is not defined, we can drop the corresponding term or assign 0 to undefined cases.

### A.6 Maximum Likelihood Computation for a Tree

Using tree traversals to lower complexity is well established (e.g. (7)) and we give details here for completeness. First we look at the cell lineage tree. Let *j* be a mutated locus of type *XY*, we want to find the likelihood of attaching *j* to each edge. Let *u* be a node in the tree and denote by *C*(*u*) the set of leaves (i.e. observed cells) below this node, then the likelihood that *j* occurs just above *u* is

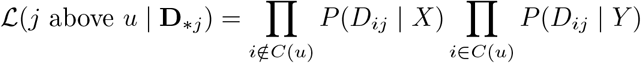

This method requires *N −* 1 arithmetic operations for all 2*N −* 1 nodes and *M* mutated loci, so that the time complexity is *O*(*N* ^2^*M*). To simplify the calculation, first note that when *u* is a leaf, *C*(*u*) is constant regardless of tree structure, in which case ℒ(*j* above *u* | **D**_∗*j*_) is the same for all trees. This halves the number of nodes that need recalculation at each step. When *u* is internal, we can calculate the likelihoods at *u* recursively from corresponding values of its children. Denote the two children of *u* by *v* and *w*, then *C*(*v*) ∩ *C*(*w*) = ∅ and *C*(*v*) ∪ *C*(*w*) = *C*(*u*). It follows that

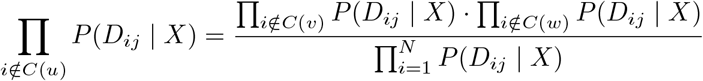

and

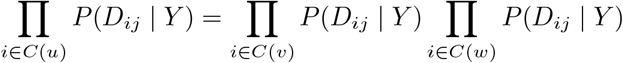

Multiplying these two terms, we get

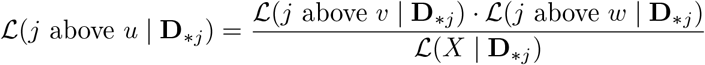

Here, the denominator ℒ(*X*|**D**_∗*j*_) is the likelihood of mutation *j* not occurring at all (so that all cells have the genotype *X*). Therefore, if we use reverse depth-first traversal, we only need 2 arithmetic operations for each node.

In addition, we can define *λ*(*j, u*) as the likelihood ratio between [*j* attached above *u*] and [*j* not occurring]

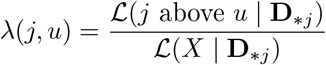

Since the denominator is constant, the arguments of maxima is the same for *λ* and ℒ, i.e.

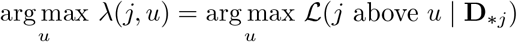

which means we can find the best attachment location with *λ*(*j, u*) as well. In this case we have

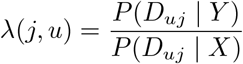

if *u* is a leaf and

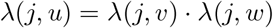

if *u* is an internal node, which further reduces the number of arithmetic operations to 1 for each node. After determining the best *u*, we can multiply *λ*(*j, u*) with the constant | (*X*|**D**_∗*j*_) to get back the likelihood. With these simplifications, it takes *O* (*NM*) time to find the highest likelihood achievable by a cell lineage tree.

For the mutation tree, we can use a similar approach that calculates the likelihood of a cell attaching to each node from the corresponding value at the parent node. Let *X*_1_*Y*_1_, …, *X*_*M*_ *Y*_*M*_ be the mutation types. Let *u* be a node in the mutation tree, *i* be an observed cell and *A*(*u*) be the set of ancestors of *u* (i.e. mutations preceding *u*), including *u* itself. Then the likelihood of attaching *i* to *u* is

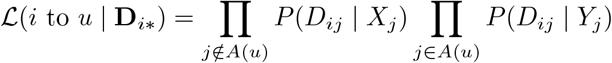

As long as *u* is not the root, it represents a mutation and has a parent node *p*(*u*), and we have *A*(*p*(*u*)) = *A*(*u*)|{*u*}. It follows that

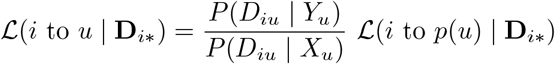

Since *P* (*D*_*iu*_|*Y*_*u*_)*/P* (*D*_*iu*_|*X*_*u*_) is independent on the tree structure, we can calculate it in advance, so that only 1 arithmetic operation remains. In case of *u* being the root, we have *A*(*u*) = ∅, and the likelihood simplifies to

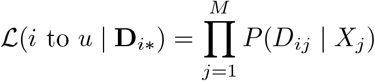

which can also be calculated in advance. Therefore, the time complexity of finding the best achievable likelihood for a mutation tree is again *O*(*NM*).

For our simple tree moves, calculating the tree likelihood for every possible attachment point can be made even more efficient. For example, if we prune and reattach subtrees of cell lineage trees, the likelihood ℒ(*j* above *u*|**D**_∗*j*_) changes only for the attachment node and its ancestors. For the rest of the tree the likelihood stays unchanged. We can thus calculate the likelihoods for each node of the subtree and the main tree in advance and only adjust it locally whenever is needed to find the optimal reattachment point.

In Figure A.3 we see that for the displayed number of cells and mutations, the runtime of the whole SCITE-RNA pipeline scales well.

### A.7 Comparison with existing methods based on clustering and bootstrapping

Both SClineager and DENDRO use hierarchical clustering to assign cells to clones and infer a cell lineage tree. To enable a direct comparison, we applied the same clustering approach to the predicted variant allele frequencies derived from the optimal likelihood SCITE-RNA tree, using the true number of clones as input for all methods. As shown in Figure A.1, SCITE-RNA with clustering consistently outperforms DENDRO and SClineager in terms of the Path Difference across all simulated settings.

The predicted variant allele frequencies were calculated by averaging the observed frequencies per clone. While our method with clustering demonstrates superior performance for datasets with few clones (Fig. A.2), it is worth noting that the predicted genotypes from SCITE-RNA can also be used directly for more accurate variant allele frequency estimation (as in Fig. 4).

For assessing the quality of clone assignments, we use the adjusted Rand index (6; 22), which quantifies the similarity between the predicted and true clusterings while correcting for chance agreement. However, the adjusted Rand index and averaging of observed variant allele frequencies per clone are not meaningful in settings where each cell represents its own clone, such as with 50 clones and 50 cells, and so are not displayed (Fig. A.2).

Bootstrapping provides a means of quantifying uncertainty in the tree structure. From a sample of 1,000 bootstrapped trees, we generated a consensus tree using `dendropy.consensus` and `resolve_polytomies` from the DendroPy package (29) (version 5.0.6), with a minimum clade frequency of 0.01. This procedure ensured that each cell was placed in the consensus tree, even when support was low. As illustrated in Figure A.1, the consensus tree more closely resembles the ground truth than individual trees without bootstrapping for the case of five clones. For more complex clonal structures, however, the benefits of consensus tree construction are less pronounced.

